# Spikeling: a low-cost hardware implementation of a spiking neuron for neuroscience teaching and outreach

**DOI:** 10.1101/327502

**Authors:** Tom Baden, Ben James, Maxime JY Zimmermann, Phillip Bartel, Dorieke Grijseels, Leon Lagnado, Miguel Maravall

## Abstract

Understanding of how neurons encode and compute information is fundamental to our study of the brain, but opportunities for hands-on experience with neurophysiological techniques on live neurons are scarce in science education. Here, we present Spikeling, an open source £25 in silico implementation of a spiking neuron that mimics a wide range of neuronal behaviours for classroom education and public neuroscience outreach. Spikeling is based on an Arduino microcontroller running the computationally efficient Izhikevich model of a spiking neuron. The microcontroller is connected to input ports that simulate synaptic excitation or inhibition, dials controlling current injection and noise levels, a photodiode that makes Spikeling light-sensitive and an LED and speaker that allows spikes to be seen and heard. Output ports provide access to variables such as membrane potential for recording in experiments or digital signals that can be used to excite other connected Spikelings. These features allow for the intuitive exploration of the function of neurons and networks. We also report our experience of using Spikeling as a teaching tool for undergraduate and graduate neuroscience education in Nigeria and the UK.

## Introduction

Neuroscience is a major arm of modern life sciences, and many universities worldwide are now offering dedicated neuroscience undergraduate degrees [1], [2]. A fundamental aspect of these courses is understanding electrical signalling within neurons and the transmission of signals across synapses [3], as well as the experimental techniques necessary to observe these properties [4]. However, owing to budgetary constraints and logistical hurdles, few students can be afforded the opportunity to experience an electrophysiological recording of a living neuron in action, for example during an experimental class. Similarly, public understanding about the fundamentals of brain function is hampered by the lack of cheap, approachable and easy-to-use tools for neuroscience outreach aimed at illuminating how the basic machines of the brain, neurons and synapses, operate to represent information [5]. The growing public interest in areas such as artificial intelligence and the effects of neurodegeneration on an aging population make it more pressing than ever to foster public awareness and interest in basic concepts in neuroscience [6].

To support university level neuroscience teaching and public understanding of neurons, we designed “Spikeling” (Fig. 1A), a £25 electronic circuit that mimics the electrical properties of a spiking neurons by running the computationally efficient yet versatile Izhikevich model [7] in real-time. The circuit is built around an Arduino [8], an open source programmable microcontroller that has found widespread use in the teaching of engineering and the design and implementation of open source laboratory hardware [9], [10].

Following the footsteps of Mahowald and Douglas’ 1991 first complete *in silico* realisation of a spiking neuron [11], Spikeling presents a simple yet powerful model of an excitable neuron with multiple dials and input/output options to play with. It is designed to facilitate a hands-on and intuitive approach to exploring the biophysics of neurons, their operation within neuronal networks and the strategies by which they encode and process information. Spikeling can be excited and its activity recorded so as to design a variety of classical experiments similar to those that might be carried out on a biological neuron and which students learn about in textbooks [12], [13]. Here, we present a series of basic neuronal processes that are efficiently modelled using Spikeling, followed by an evaluation of our experience using the device for teaching senior undergraduate and MSc students in the UK and a graduate neuroscience summer school held in Nigeria. Spikeling should be a useful tool in educating students of neuroscience and psychology, as well as students of engineering and computer science who are interested in the biophysics of neurons and brain function.

## MAIN

### A simple hardware implementation of a spiking neuron

Spikeling (Fig. 1) consists of an Arduino-Nano microcontroller, a custom-printed circuit board, and a small number of standard electronic components (see Bill of Materials, BOM). Assembly takes between 20 minutes and 2 hours, depending on previous experience with soldering and assembling circuit boards (see Spikeling manual). Spikeling features large contacts and ample component spacing to facilitate soldering for beginners. The functional properties of Spikeling can be modified by software within the Arduino integrated development environment (IDE).

Upon current injection, Spikeling begins to fire, with each spike translating into an audible “click” from a speaker. In tandem, membrane potential is continuously tracked by the brightness of a light-emitting-diode (LED). To mimic different types of neurons, Spikeling features a “mode button” for switching between different pre-programmed model behaviours (e.g. regular spiking, fast spiking, bursting etc.). These can also be modified in the code provided.

**Figure 1.**
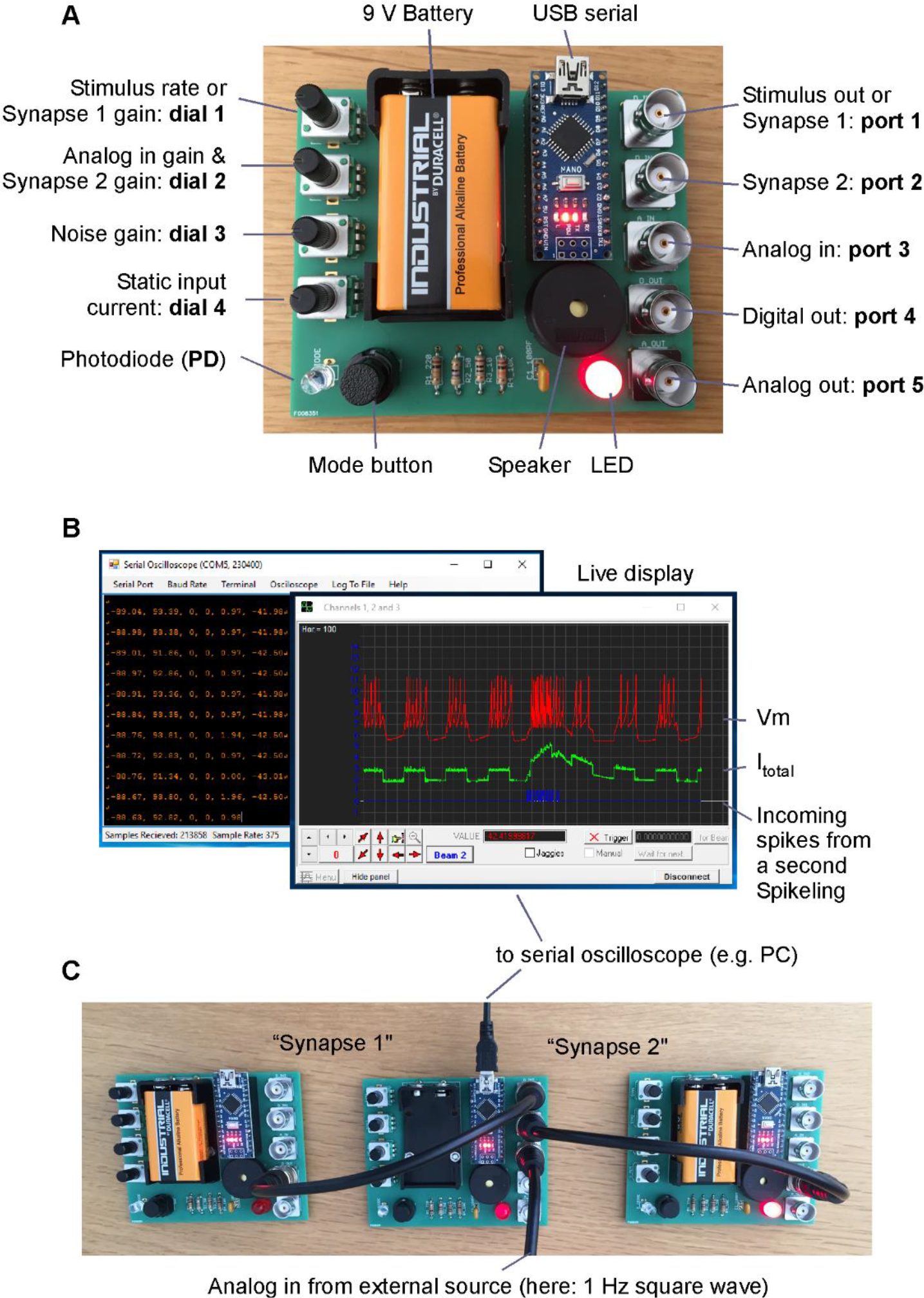
Basic hardware and software. **A**, Fully assembled Spikeling board. **B**, Screenshots of the Serial Oscilloscope software [14] used displaying Spikeling activity of the network in (C). **C**, Three Spikelings connected into a simple network.

For inputs, Spikeling (Fig. 1A, Supplementary Figure S1) has three Bayonet Neill-Concelman (BNC) ports: Two are “input synapses” that each respond to 5V transistor-transistor-logic (TTL) pulses (ports 1,2) such as the “spike output” of a second unit. Thus, Spikelings can also be connected into simple neuronal networks (Fig. 1B, C). A third BNC input connection (port 3) is an analog-in port that can be driven with a stand-alone stimulus generator or by a computer with a suitable output port. The gain and sign of all inputs can be continuously set with rotary encoder knobs (dials 1 & 2 - with dial 2 controlling both analog-in and synapse 2 gain). One aim in the design of Spikeling was to also teach how neurons encode a sensory stimulus so an on-board photodiode allows Spikeling to sense light. A light stimulus can be delivered externally (e.g. using a torch), or via an LED driven by a programmable on-board pulse generator. To mimic the “noisiness” of biological neurons in intact neural circuits, a knob is provided to add variable amounts of membrane noise to the simulation (dial 3) while a final knob controls a static input current to set resting membrane voltage (dial 4).

For outputs, Spikeling features digital (port 4) and analog (port 5) BNC connections that can be used to visualise the “membrane voltage” output on an external oscilloscope or to drive another Spikeling. Alternatively, the modelled membrane potential and several key internal processes (e.g. different current sources, input spikes etc.) can be directly read out through the USB-based serial port for live display on a computer screen and data logging (Fig. 1B). We also provide Matlab (Mathsworks) scripts for basic data visualisation and analysis. Finally, the system can be powered through the universal serial bus (USB) port or by a 9V battery.

### Simulating neuronal activity

In an informal setting, Spikeling can be explored in a playful manner simply by (i) depolarising or hyperpolarising the neuron via the input current (dial 4), (ii) dialling up the membrane noise (dial 3, Fig. 2A) or (iii) manual stimulation of the photodiode with a torch (Fig. 2B, SVideo 1). In each case, elicited spike activity can be intuitively tracked by audible clicks coupled to flashes of the onboard LED. In parallel, membrane potential and input current can be tracked live on a PC screen through a serial plotter such as the openly available “Serial oscilloscope” [14] (Fig. 1B). In this setup, Spikeling can be used to explore basic concepts in neuronal coding. For example, holding a torch over the photodiode initially elicits a burst of spikes that gradually slows down if the light is held in place, thereby mimicking a slowly adapting “light-on” responsive neuron (Fig. 2B, left). The same experiment with Spikeling set to mode 2 (toggled via the onboard button) will reveal a rapidly adapting rebound burst of spikes upon removing the light, thereby mimicking a transient light-off responsive neuron (Fig. 2B, center). Next, mode 3 mimics a sustained light-off driven neuron with an elevated basal spike rate (Fig. 2B, right, cf. SVideo 2). In total, Spikeling is pre-programmed with 5 modes (Supplementary Figure 2). These can easily be modified or extended by the user in the Arduino code provided.

**Figure 2.**
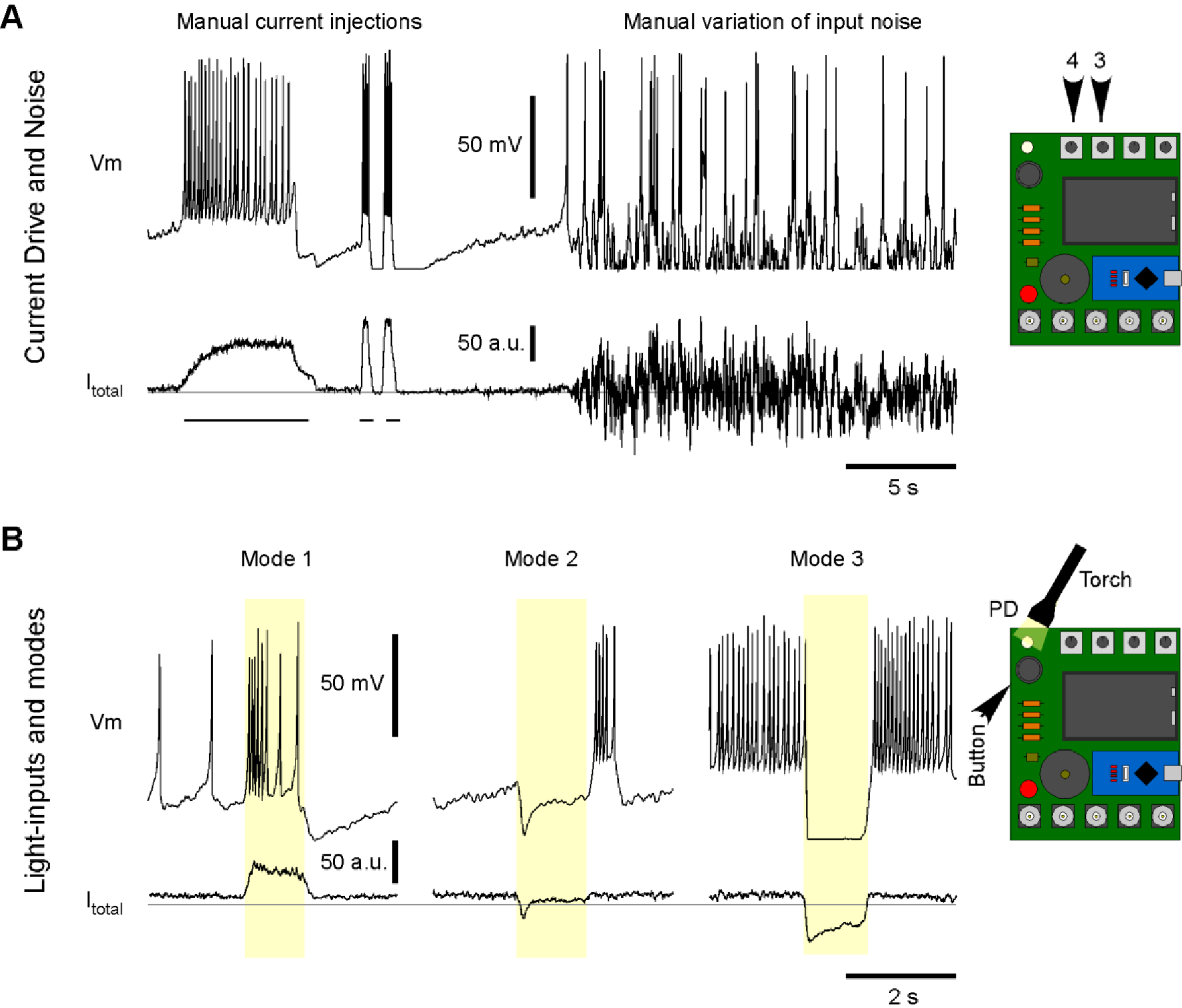
Manual exploration of Spikeling functions. **A**, Example recording of Spikeling membrane potential (top) and current (bottom) during manual manipulations of the input current dial (4) to depolarise the neuron (left) and following the addition of a noise current (dial 3, right). **B**, Example light responses in modes 13 (left to right, toggled by the button) to manual photodiode (PD) stimulation with a torch. The grey horizontal lines indicate I_total_ = 0.

For more formal experimentation, Spikeling can be driven in a temporally precise manner via the analog-in port or a regularly pulsed light source mounted over the photodiode (SVideo 3). As a stimulus, port 1 (synapse 1/ stimulus out) can be flexibly reconfigured into a digital stimulus generator. Alternatively, an external 0-5V analog stimulus generator can be connected (not shown). At default settings, this port will continuously generate 0-5V pulses at 50% duty cycle, with the stimulation rate being controlled through dial 1. Accordingly, simply connecting port 1 (stimulus out) to port 3 (analog-in) allows for simple, time precise stimulation of the model neuron.

The millisecond time precision achieved in this way can then be exploited to study neuronal function in further detail. For example, at default settings (see Spikeling manual) the stimulator directly coupled to the analog-in port drives a highly stereotyped spike train upon repeated stimulation (Fig. 3A, left), as further elaborated in the raster plot (Fig. 3A, right, see also Supplementary Figure S2). From here, systematic variation of the analog-in gain (dial 2) can be used to drive Spikeling with different amplitude current steps, for example to build amplitude tuning functions for spike rate, latency or first-spike time-precision (Fig. 3B).

Next, rather than delivering port 1’s square-pulse drive via analog-in, the user can instead drive an LED from the same port. In this way, positioning the LED above the photodiode (e.g. via the 3D-printable adapter provided, or a custom paper tube) allows for temporally precise driving of Spikeling via light (Fig. 3C). Adding noise to this simulation allows exploring how the addition of noise initially distorts spike timings before affecting rates (Fig. 3D).

**Figure 3.**
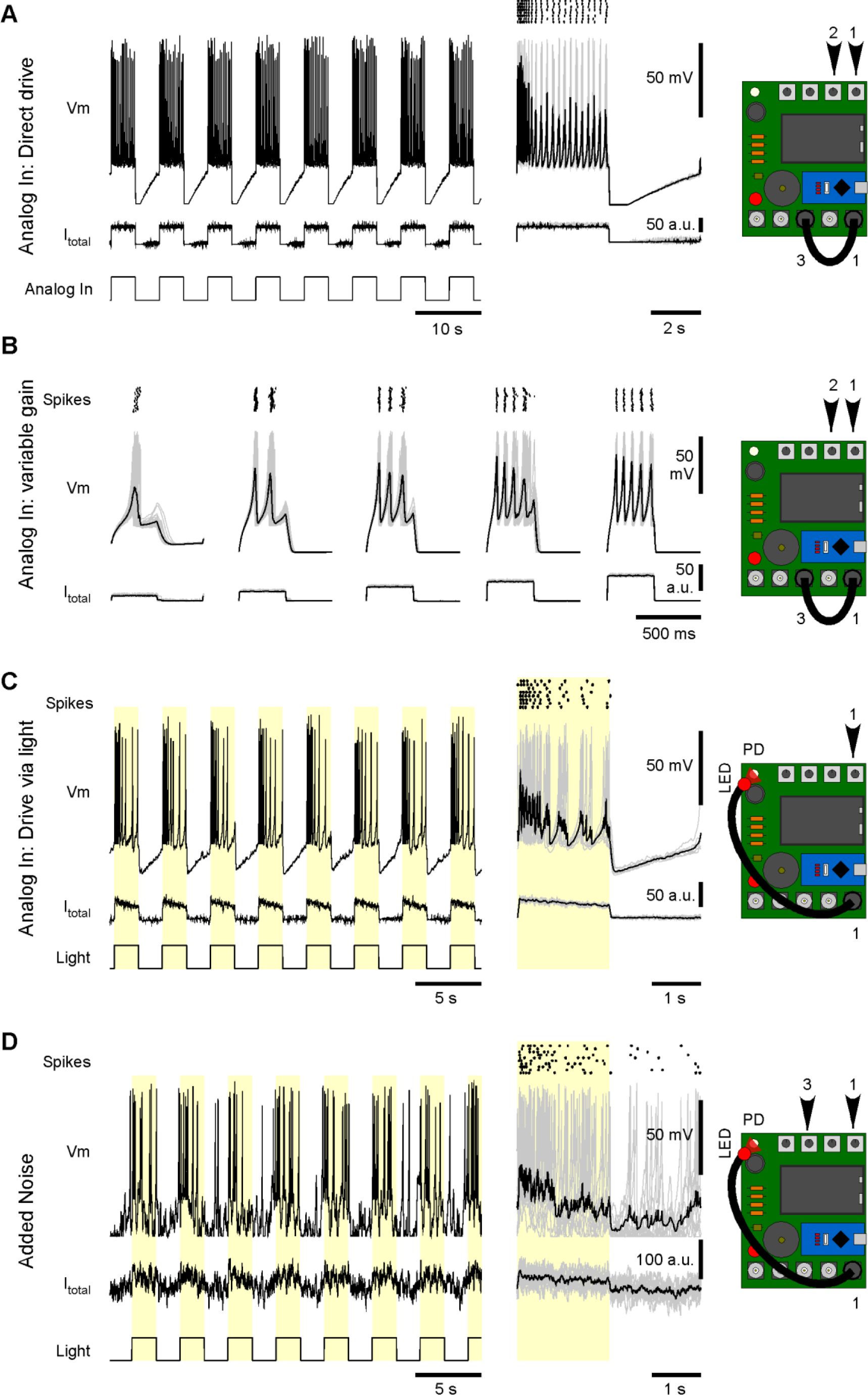
Basic stimulus-driven functions. **A**, Example recording of Spikeling in Mode 1 driven by the internal stimulator (port 1) via the Analog In connector (port 3) as indicated. Gain and stimulus rate are controlled on dials 2 and 1, respectively. Right: stimulus aligned response segments (grey) and average (black) as well as spike raster plot. **B**, as (A, right), with varying input gain to probe amplitude tuning. Note systematic effects on spike number, rate, time latency and time precision. **C**, As (A), but this time driving Spikeling via an LED attached to the stimulus port stimulating the photodiode. Note different waveforms of input current and consequences on the elicited spike pattern compared to (A). **D**, as (C), with addition of current noise (dial 3). Note distortion of spike timings, while the number of spike triggered remains approximately constant.

Similarly, the experimenter could vary the rate of stimulation to probe the intrinsic frequency tuning of a neuron (dial 1, not shown). At faster stimulus rates, Spikeling can be set to occasionally “miss” individual current steps and instead adopt a volley code [15] for event timing (Fig. 4A). In this configuration, Spikeling continues to phase-lock to the stimulus, as summarised in the event-aligned plot to the right. Note that even though spikes frequently fail, the subthreshold potential continues to reliably track the stimulus. From here, the static input current (dial 4) and noise (dial 3) can be tweaked to put the system into stochastic resonance [16], [17]: In this situation, counterintuitively, the addition of noise is beneficial to the code (Fig. 4B). In the example shown, the “generator potential” (the noise-free stimulus driven membrane voltage fluctuations) is itself insufficient to elicit any spikes. As a result, the neuron fails to encode the stimulus at the level of its spike output (Fig. 4B, left). However, addition of noise occasionally takes the combined generator potential and noise above spike threshold (Fig. 4B, middle). Importantly, the probability of this threshold crossing is higher during a depolarising phase of the generator. As a result, the system now does elicit spikes which, depending on the noise level chosen, reliably phase-lock to the stimulus (Fig. 4B, right). Such stochastic resonance can be used e.g. by sensory systems to deal with noisy inputs - summing across the spike output from many such resonating neurons can then reconstruct the original stimulus with high fidelity [18], [19].

**Figure 4.**
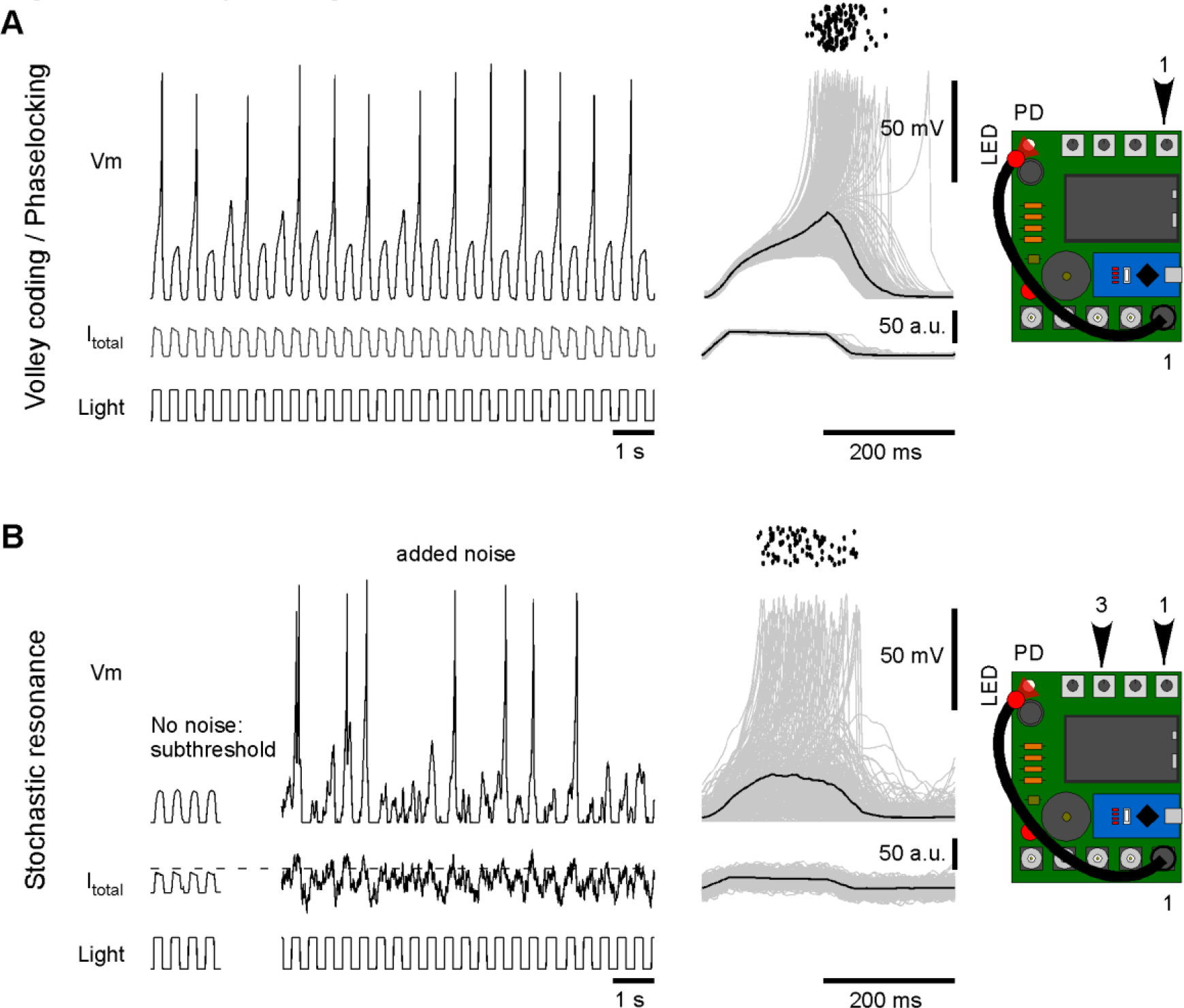
Volley coding and stochastic resonance. **A**, By varying the stimulus rate, Spikeling can be set-up to “miss” individual stimulus cycles at the level of the spike output (left). However, when elicited, spikes remain phase-locked to the stimulus (right). **B**, Example of stochastic resonance: as (A), with neuron hyperpolarised just enough to prevent all spikes (left). Now, addition of membrane noise occasionally elicits spikes (middle), which again are phase-locked to the stimulus (right). Dotted line indicates approximate spike threshold.

Next, two or more Spikelings can be connected into a network via BNC cables (SVideo 4). For this, the digital-out connector (port 4) of one unit is connected to one of two “synapse-in” connectors (e.g. port 2) on another unit. Synaptic gain can then be controlled using a rotary encoder (here: dial 2) to vary the efficacy and sign of the coupling, thus mimicking excitatory or inhibitory connections (Fig. 5A). Two reciprocally connected units can then be used to set up a basic central pattern generator [20], [21] (Fig. 5B).

**Figure 5.**
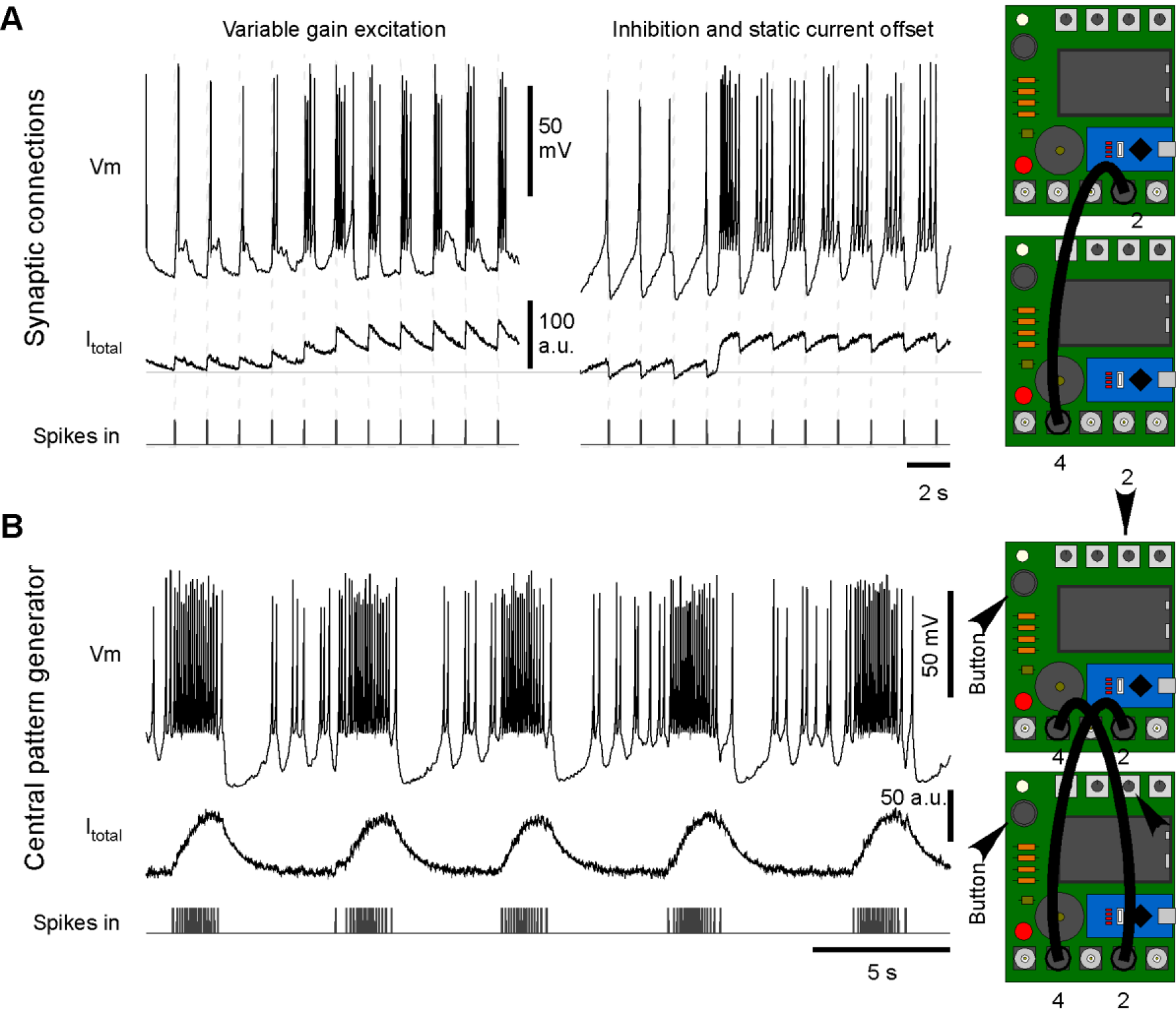
Synaptic Networks. **A**, Two or more Spikelings can be connected to form synaptic connections, as indicated. Left: Excitatory synaptic connection with synaptic gain gradually increased by hand over time (dial 2). Right: Inhibitory connection at two different depolarisation states (dial 4). **B**, Example of a 2-neuron central pattern generator (CPG). The two Spikelings are set to mode 2 and wired to mutually excite each other. In each case, all traces display the activity and incoming spikes of the top-most Spikeling.

Spikeling can also be used to explore neuronal function more systematically, for example by estimating the linear filter that underlies its photo-response in a given mode [22]. This is a fundamental approach in computational and sensory neuroscience, and the calculation of the linear filter is based on recording a neuron’s response to a “noise stimulus” for several minutes. Subsequent reverse correlation of the elicited spike-or subthreshold activity against the original stimulus then allows calculating the average stimulus that drove a response in the neuron: the linear filter, sometimes also referred to as “time-reversed impulse response” or “response kernel”. Reverse correlation to spikes is the more common calculation, when the linear filter is also termed the “spike-triggered average” or STA [23]. To explore this concept, Spikeling’s stimulus port (1) can be set to generate binary noise at e.g. 50 Hz via a flag in the Arduino code (see Spikeling manual). In this configuration, the photodiode can be stimulated by this noise stimulus via an LED as before (Fig. 6A, cf. Fig 3C), thereby driving spikes and subthreshold oscillations. The linear filters of a mode 1 Spikeling (“slow”) reveal a clear biphasic (band pass) stimulus dependence at the level of spikes, but a monophasic dependence (low pass) at the level of subthreshold activity (Fig. 6B, black). In comparison, the same mode 1 neuron retuned to use a rapidly adapting photodiode-driven current (“fast”) gives a triphasic stimulus dependence at the level of spikes and a biphasic dependence at the level of the subthreshold generator (Fig. 6B, red).

**Figure 6.**
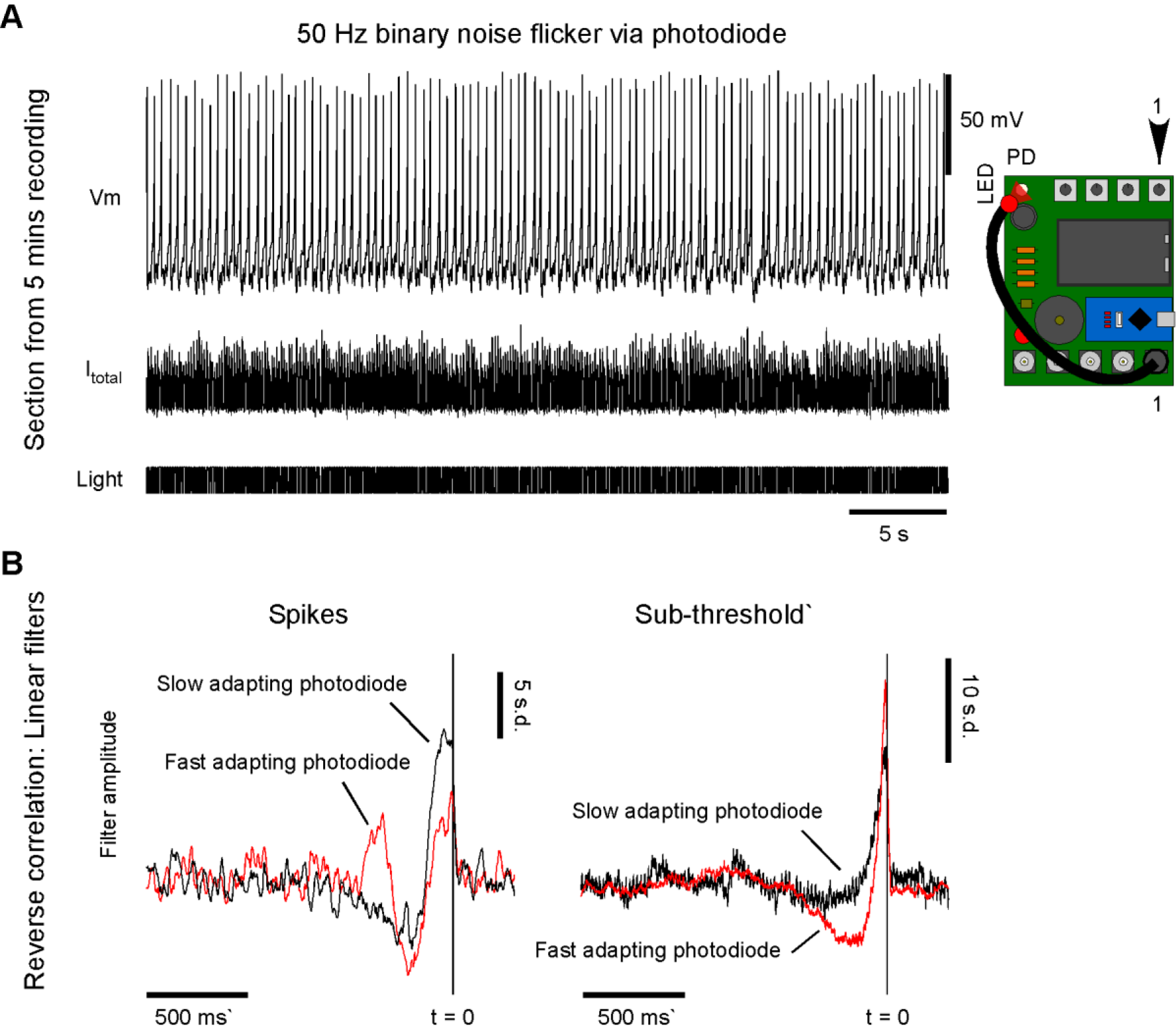
Estimating linear filters by reverse correlation. **A**, Via the Arduino code, the stimulus port can be set to deliver 50 Hz binary noise, here used to drive the photodiode via an LED (cf. Fig 3C). Current and spike pattern elicited by this stimulus. **B**, linear filters of a slow (black) and a fast (red) photo-adapting mode 1 neuron estimated at the level of spikes (left) and subthreshold membrane potential (right).

Taken together, Spikeling can be used in a variety of classroom and demonstration scenarios, ranging from simple observations of changes in spike rates upon stimulation to advanced concepts in neuronal computation and analysis.

An example set of Spikeling-based classroom exercises is provided (see Spikeling manual). From here, advanced users can easily re-programme the Arduino code to implement or fine tune further functionalities as required. The entire project, including all code, hardware design, bill of materials and detailed build instructions are available online for anyone to freely view and modify (https://github.com/BadenLab/Spikeling and https://badenlab.org/resources/).

### Spikeling in the classroom

We evaluated the utility of Spikeling in two classroom scenarios: (i) as a 2-day section within a 3-week intensive neuroscience summer school held at Gombe State University, Nigeria by TReND in Africa [24] and (ii) as part of an 18 lecture module on *Sensory function and computation* delivered for 3^rd^ year undergraduate and MSc neuroscience students at the University of Sussex, UK. We report on each experience in turn.

At Gombe State University, Nigeria, we ran two identical 2-day sessions for a total of 18 Africa-based biomedical graduate students (9 at a time) as part of the 7^th^ TReND/ISN school on Insect Neuroscience and *Drosophila* Neurogenetics [24]. None of the students had much experience with neuronal computation or electrophysiological techniques, although most had covered basic concepts in neuroscience such as action potential generation in their undergraduate degrees. We introduced Spikeling in three steps. First, we held a 1-hour lecture where a single Spikeling was connected to a computer with the serial oscilloscope output being projected live to the wall. In parallel, a whiteboard was used for explanations and discussions. From here, we combined a general explanation of concepts in neuronal computation on the board (for example, rate-versus time-coding, sub-threshold integration, phase-locking etc.) and then demonstrated each phenomenon in front of the class using Spikeling. Based on feedback after the class, this was perceived as a very engaging and effective method for introducing concepts in neuronal coding. Next, we moved on to assembling Spikelings from bags of pre-compiled parts. For this, every student was provided with the printed circuit board, the electronic components and a soldering iron and taken through the assembly process by two instructors. After 2-3 hours, every student had successfully assembled a working unit, despite most not having had any experience with soldering or electronic circuit logic. In a third step, each student was then provided with the serial oscilloscope software as well as the exercise documentand asked to sequentially work through a set of pre-designed exercises (see Spikeling manual) in their own time, with faculty being available to help as required. Following the course, all students kept their Spikeling to facilitate their own teaching at their host institutions in 7 different African countries (Nigeria, Malawi, Sudan, Egypt, Kenya, Zambia, Burkina Faso).

At the University of Sussex, UK, we introduced pre-assembled Spikelings as part of 3 sets of 3-hour workshops that each ran twice to accommodate 26 students in groups of 13. For this, we used a PC lab where each student had their own Spikeling and PC with Arduino, Serial Oscilloscope and Matlab preinstalled. The first session began with a 20 mins presentation of basic concepts in neuronal modelling and electronics followed by a conceptual comparison between the biophysically realistic yet computationally heavy Hodgkin Huxley model [25], [26] and the much lighter phenomenological Izhikevich model [7] implemented in Spikeling. Next, we projected the serial oscilloscope screen of one Spikeling connected to the lecturer’s laptop to the wall. This allowed easy, live demonstrations of some Spikeling functions, such as the photo-response or the use of different modes. From here, we asked students to connect and set up their own units on their PCs and to start exploring “how to best drive spikes” using their mobile phone torches. Students quickly realised that simply holding the light above the photodiode ceases to be effective after a few hundred milliseconds, while repeatedly moving the light over the photodiode reliably elicits bursts of spikes. In this way, students could intuitively explore basic concepts in time coding. Afterwards, we brought everyone back to the same page by demonstrating those key ideas on the Spikeling output projected to the wall. We then showed students how to use the stimulator, what the dials do, and how to log data on the serial oscilloscope. We also showed how to load and display their data using prewritten Matlab routines (see Supplementary Materials). From this point, we asked students to themselves quantify a neuron’s amplitude tuning at the level of instantaneous spike rate and first-spike latency to compare the two, again followed by an in-class demonstration and discussion afterwards. In this way, we moved through the majority of Spikeling functions described in this paper over the course of 3 workshops.

Taken together, Spikeling allowed students to explore a number of fundamental aspects in sensory neuroscience, including analog and digital coding, detection of signals above noise, the functional consequences of adaptation, and the variety of temporal filters that neurons implement. The concepts acquired, as tested with take-home problem sets, dovetailed with lecture content covering rate and time coding, feature selectivity and tuning diversity, and adaptation. Students reported that the Spikeling work helped them to develop a more intuitive grasp of these central ideas in sensory systems neuroscience.

## Discussion

With modern systems neuroscience increasingly moving into the area of big data where the activity of 1,000s of neurons can be routinely recorded across a wide range of neuronal circuits [27]-[33], a deep understanding of how neurons encode and compute information is fundamental. These concepts need to be taught not just to students of the biological sciences but also to students of psychology as well as engineers and computer scientists interested in theoretical and computational neuroscience, artificial intelligence and robotics [4]. However, concepts in neuronal coding and computation can be unintuitive to grasp or “dry” in lectures, while classroom electrophysiology on live biological specimens can be technically challenging and costly to set up [3]. As a result, many students in these disciplines graduate without ever having had the opportunity to experience and control neuronal activity in hands-on experiments. Indeed, in many parts of the world, systems neuroscience is only a rather peripheral aspect of neuroscience curricula, if present at all, while the cross-over of neuroscience into engineering and informatics often jumps immediately into discussions of networks based on units that are greatly simplified versions of biological neurons.

Spikeling is intended to help ameliorate some of these issues by allowing students to carry out experiments in the same general fashion as classical electrophysiologists but without the amplifiers, filters, manipulators, stimulus generators and other equipment normally required. Its low cost makes it widely affordable, and once assembled, it can be used for teaching for many years without additional investment. It should also be immediately approachable to students of engineering and informatics who can explore the electrical properties of neurons and the code used to model these as well as carry out experiments illustrating basic concepts in theoretical and computational neuroscience [23]. By allowing students to interact physically with the device, e.g. by providing actual sensory inputs, Spikeling can help build an intuitive grasp of neuronal computations beyond that provided by pure computer simulation of neurons.

Other recent efforts have also recognised the need for more intuitive hardware models of spiking neurons, most notably the Neurotinker^®^ initiative [34] who release NeuroBytes^®^. In this case, the design is geared towards schools and the general public to facilitate playful exploration of neuronal function and, in particular, networks. Another initiative aiming to build microcontroller-based neurons is Spikee [35].

With time, we hope that others may take up our basic design and build upon it, for example by providing inputs to other sensory modalities such as touch or sound or by changing the Arduino code to implement new functions or simulate neurons with different tuning properties. Spikeling is available on a share-alike open license, prompting any modifications of the original code to be freely re-shared for everyone to use. We aim to keep these efforts centralised on the Spikeling GitHub, or link to new repositories as they arise to gradually build a community of users and contributors.

## Acknowledgements

We thank Peter Reed and Chen Qian for technical advice and Ihab Riad for helping with in-class Spikeling assembly and teaching in Nigeria. The authors would also like to acknowledge support from the FENS-Kavli Network of Excellence. Funding was provided by the European Research Council (ERC-StG “NeuroVisEco” 677687 to TB), Marie Sklodowska-Curie European Training network “Switchboard” to TB (Switchboard receives funding from the European Union’s Horizon 2020 research and innovation programme under the Marie Sklodowska-Curie grant agreement No. 674901), The Deutsche Forschungsgemeinschaft (DFG: BA 5283/1-1 to TB), The Medical Research Council (TB, MC_PC_15071) and the Leverhulme Trust (PLP_AAF_20171024 to TB). In addition, we would like to thank the International Society of Neurochemistry (ISN), the Company of Biologists (CoB), The Cambridge Alborada Fund and hhmi Janelia Research Campus for supporting our work in Nigeria.

## Author contributions

TB conceived of, designed and implemented Spikeling. The Matlab pre-processing scripts were written by BJ, with modifications by TB. Spikelings for UK teaching were assembled and tested by MYZ, PB and DG. All authors contributed to in-class teaching and evaluation in the UK. TB taught the course in Nigeria. The paper was written by TB with help from LL and MM and inputs from all authors.

## Data availability

All Hardware instructions, code, manuals and example data are freely available at: https://github.com/BadenLab/Spikeling and https://badenlab.org/resources/.

---

**Supplementary Figure 1.**
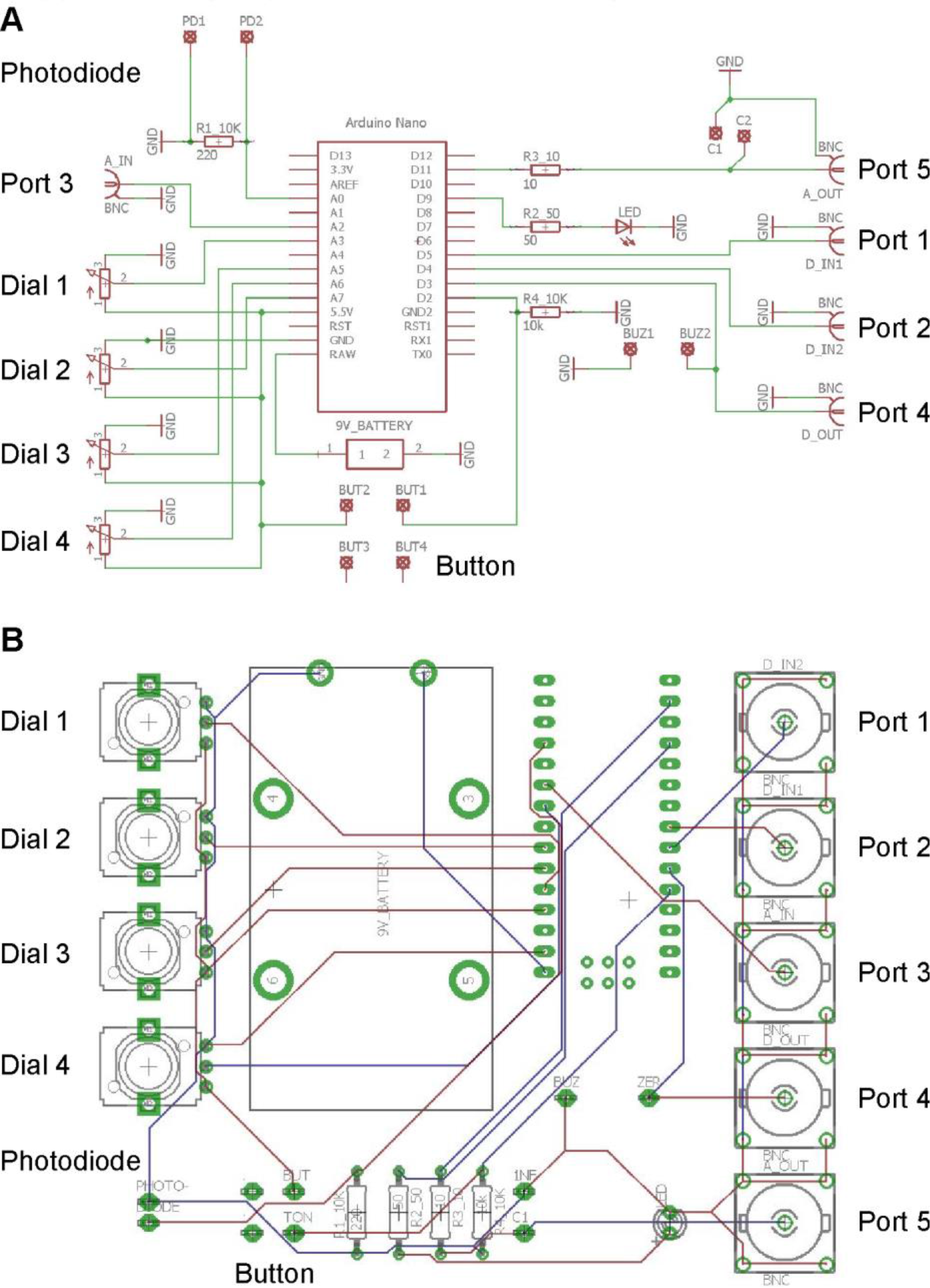
Circuit and PCB layout. **A,** Wiring diagram of Spikeling. **B**, PCB Layout.

**Supplementary Figure 2.**
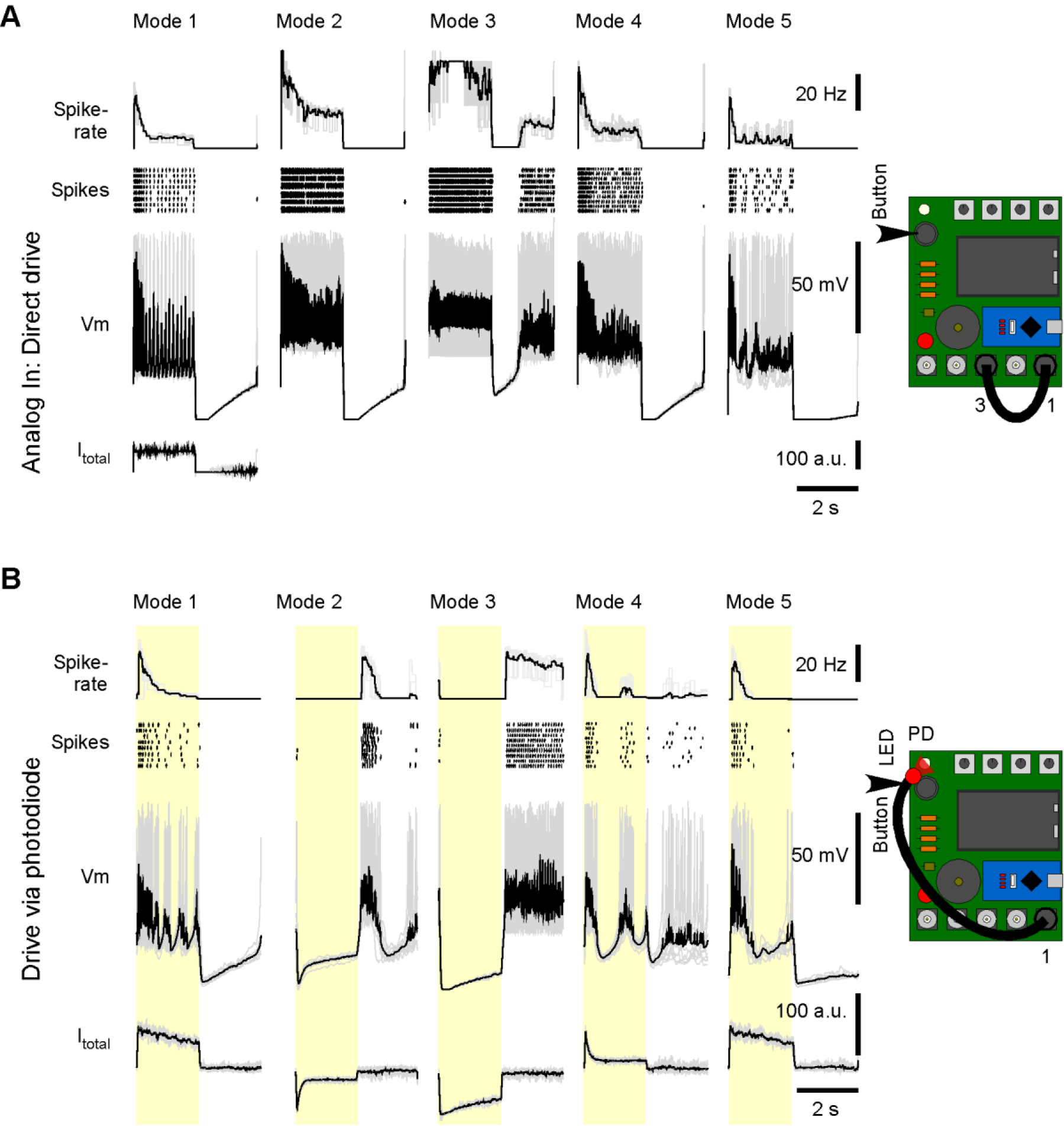
Mode Overview. **A,B,** All 5 pre-programmed Spikeling modes responding to current (A) and light steps (B). Additional modes can be easily added in the Arduino code (see Spikeling manual).

**Supplementary Video 1: Basic functions**

**Supplementary Video 2: Modes Supplementary**

**Video 3: Stimulus generator Supplementary**

**Video 4: Synaptic Networks**

**Supplementary data files provided**:

– Spikeling Manual including assembly and example exercises
– Bill of Materials (BOM)
– PCB layout files (Eagle)
– Arduino code (x2) for Spikeling
– Matlab code (x2) for basic data analysis and visualisation
– OpenSCAD and surface-tessilation (stl) files for 3D-printable LED-mounting adapter
– Example logged data (csv)

